# Longitudinal Salivary Immunophenotyping Reveals Distinct Cellular Signatures of Periodontal Disease Activity and Resolution

**DOI:** 10.64898/2026.07.01.735878

**Authors:** Raza Ali Naqvi, Matt Tokarski, Kristofer Ceredon, Joseph Gluck, Sarah Elshourbagy, Laura Popa, Lana Dalbah, Michael L. Schmerman, Joel L. Schwartz, Salvador Nares, Afsar R. Naqvi

## Abstract

**Aim:** To investigate whether salivary immune cell profiling can serve as a non-invasive approach to monitor periodontal disease activity and therapeutic response by characterizing innate and adaptive immune cell dynamics in periodontitis.

**Materials and Methods:** This longitudinal study included systemically healthy adults with periodontitis and healthy controls. Periodontal parameters (PPD, BOP, plaque/calculus, and radiographic bone loss) were recorded by calibrated examiners (κ=0.85) following established criteria. Stimulated saliva and gingival biopsies were collected before and 4–6 weeks after non-surgical periodontal therapy (NSPT), and from healthy controls. Multiparametric flow cytometry was used to characterize myeloid and lymphoid cell populations and polarization markers. Bacterial transcripts and host inflammatory markers were assessed by qRT-PCR. Statistical analyses were performed using one-way ANOVA.

**Results:** Periodontitis subjects exhibited significantly elevated salivary bacterial transcripts, which decreased but did not normalize following NSPT. Both myeloid and lymphoid immune cell populations increased in periodontitis compared with healthy controls and declined after therapy. This was accompanied by a pronounced pro-inflammatory shift with elevated IFN-γ–producing macrophages, dendritic cells, Th1/Th17 cells, and B cells, including the novel identification of IFN-γ–producing B cells in saliva and mirrors the gingival immune cell profiles. In contrast, anti-inflammatory populations (IL-10–producing myeloid cells, Tr1 cells, and regulatory B cells) were reduced in disease and partially restored following NSPT.

**Conclusions:** Salivary immunophenotyping non-invasively monitors PD activity and therapeutic response by capturing dynamic immune changes that reflect gingival signatures and track post-therapy resolution.

**Clinical Relevance:** *Scientific rationale:* Salivary immune profiling offers a real-time, non-invasive tool for assessing periodontal disease status and treatment outcomes, with potential applications in precision diagnostics and personalized periodontal care.

*Principle findings:* Periodontitis was associated with increased salivary bacterial burden and a marked pro-inflammatory immune profile involving both innate and adaptive immune cells, including newly identified IFN-γ–producing B cells. Non-surgical periodontal therapy partially restored anti-inflammatory immune responses and reduced inflammatory cell populations, supporting salivary immunophenotyping as a promising non-invasive biomarker approach for monitoring disease activity and treatment response.

*Practical implications:* Salivary immune cell profiling could serve as a simple, non-invasive tool to monitor periodontal disease activity and response to therapy in clinical practice. Identification of specific inflammatory cell subsets may also aid in developing personalized diagnostic and therapeutic strategies for periodontitis.

## 1. Introduction

Periodontitis is a chronic, multifactorial inflammatory disease with a significant global health burden (O’Dwyer et al., 2023, Wang et al., 2025). Longitudinal monitoring across states of periodontal health, disease, and post-therapy is essential for understanding disease progression and treatment outcomes (Sanz-Martín et al., 2019, Loos et al., 2020). However, conventional clinical and radiographic assessments primarily reflect cumulative tissue destruction and fail to capture real-time disease activity (Tonetti, et al., 2018, Liukkonen et al., 2018). This limitation highlights the need for non-invasive approaches capable of dynamically monitoring periodontal status.

Saliva has emerged as a promising non-invasive diagnostic medium for monitoring periodontal health. It reflects both oral/periodontal status and systemic physiology and can be collected repeatedly with minimal discomfort (Ji et al., 2015, Isho et al., 2020, Surdu et al., 2025,). To date, salivary diagnostics have largely focused on soluble biomarkers, including pro-inflammatory cytokines, matrix metalloproteinases, acute-phase proteins, and microbial components, which provide insight into inflammation, tissue degradation, and dysbiosis (Miller et al., 2010, Isho et al., 2020).

Despite these advances, soluble biomarkers alone may not fully capture the complexity of periodontitis, which is fundamentally driven by dysregulated host immune responses. Therefore, direct assessment of immune cell populations may provide a more mechanistic and comprehensive understanding of disease activity than measurement of downstream soluble mediators. Saliva contains an array of immune cells, including neutrophils, monocytes, macrophages (Mφ), dendritic cells, and lymphocytes, derived from periodontal tissues and oral mucosa (Vidovićet al., 2012; Choudhary et al., 2020).

These cells reflect ongoing immune processes within the periodontal microenvironment. Monitoring immune cell dynamics, such as Mφ and T cell polarization, and B cell activity, offers the potential to capture real-time immunological changes associated with disease initiation, progression, and resolution. Compared with soluble biomarkers, immune cell profiling provides higher-resolution insight into host immune status and may improve discrimination between periodontal health, disease, and therapeutic response (Choudhary et al., 2020, Theda et al., 2018).

However, despite the recognized importance of immune mechanisms in periodontitis, there remains limited knowledge regarding the composition, phenotype, and clinical utility of salivary immune cells for longitudinal disease monitoring. In particular, few studies have systematically evaluated salivary immune cell profiles across different clinical states, including periodontal health, untreated disease, and post-therapy conditions (Miller et al., 2010, Isho et al., 2020, Teles et al., 2024).

In this study, we investigate salivary immune cell profiles as potential biomarkers for periodontal disease monitoring. Using multiparametric flow cytometry, we characterize myeloid and lymphoid cell populations in saliva from healthy individuals and compare findings to subjects with periodontitis and responses to periodontal therapy. By examining immune cell dynamics across these clinical states, we aim to determine whether salivary immune profiling can provide a sensitive, non-invasive approach for distinguishing periodontal health from disease and for tracking therapeutic response.

## 2 Materials and Methods

### 2.1 Study design and population

This study was approved by the Ethics Committee at The University of Illinois Chicago, College of Dentistry (IRB Protocol# 2017-1064). Systemically healthy subjects aged 18–70 years presenting to the Postgraduate Periodontics Clinic at the College of Dentistry, from October 2023 to October 2025 were recruited for this study as we previously described (Uttamani et al., 2023, 2024). Subjects in the experimental group (n=20) exhibited stage III grade B periodontitis with probing depths ≥6 mm, clinical attachment loss ≥5 mm, bleeding on probing, and radiographic evidence of bone loss. Healthy control periodontal subjects (n=12) displayed probing depths ≤3 mm, no bleeding on probing, no evidence of clinical attachment loss, and no radiographic evidence of bone loss. Clinical parameters included periodontal probing depth (PPD), bleeding on probing (BOP), plaque/calculus, and radiographic bone loss using a PCPUNC-15 probe (Table 1). Measurements were obtained by calibrated examiners (κ=0.85). Exclusion criteria included systemic conditions affecting periodontal status (e.g., diabetes, hepatitis, renal disease, coagulation disorders, human immunodeficiency virus), smoking, antibiotic use within one month, and medications influencing gingival conditions (e.g., phenytoin, calcium channel blockers, cyclosporine).

### 2.2 Saliva and gingiva collection and processing

Saliva and gingival tissue samples were collected from periodontitis subjects before and after NSPT and from healthy controls. Detailed methods for sample collection, processing, and immune cell isolation are provided in the Supplemental Methods.

### 2.3 Wright-Giemsa Staining

Wright–Giemsa staining (Sigma-Aldrich, St. Louis, MO, USA) was performed to visualize salivary cellular components following the manufacturer’s protocol. Briefly, 0.5 mL of saliva was smeared onto a glass slide, air-dried, stained with Wright–Giemsa stain for 2 min, rinsed with deionized water, and air-dried again. Images were captured in bright-field mode using an Olympus Corporation FSX-100 microscope.

### 2.4 Flow cytometry

Saliva-or gingiva-derived cells (3×10□) were stained with fluorochrome-conjugated antibodies against CD4, CD8, CD19, CD14, CD11b, CD11c, CD56, CD66b, CCR3, CD5, and HLA-DR (all from BioLegend, San Diego, CA, USA). Intracellular staining for IL-4, IL-10, IL-17, and IFN-γ (BioLegend) was performed following fixation and permeabilization according to manufacturer’s instructions. Data (∼100,000 events/sample) were acquired using a Cytek Aurora flow cytometer (Cytek Biosciences, Fremont, CA, USA) and analyzed with Kaluza software (Beckman Coulter Life Sciences, Indianapolis, IN, USA). Multiparametric flow cytometry enabled characterization of T cell subsets (Th1, Th2, Th17, Tregs), B cells, Mφ polarization (M1/M2), and dendritic cell subsets including DC10.

### 2.5 RNA isolation and qRT-PCR

Total RNA was isolated from saliva using RNeasy kit (Qiagen). cDNA synthesis (250 ng RNA) was followed by qRT-PCR to detect bacterial transcripts (Supplementary Table 1). Gene expression was normalized using GAPDH as an endogenous control. The Ct values of three replicates were analyzed to calculate the fold change using the 2□−□^ΔΔCt^ method.

### 2.6 Statistical analysis

Data was analyzed using GraphPad Prism (GraphPad Software, Boston, MA, USA) and are presented as mean ± SD or SEM. Comparisons were performed using Student’s t-test or one-way ANOVA. Age and gender distributions were analyzed using Mann–Whitney U and McNemar χ² tests, respectively. Statistical significance was set at p<0.05.

## 3 Results

### 3.1 Elevated levels of periodontopathic bacteria transcripts in saliva of periodontitis subjects

This pilot study evaluated 32 participants to determine how non-surgical periodontal therapy (NSPT) affects pocket depth (PD), bacterial load, and immune profiles (Figure S1A). Demographic analysis showed no significant differences in age (p=0.49) or gender (p=0.55) between the cohorts (Table 1). The healthy group (mean age 49.67) maintained a mean PPD of 2.59 mm. In the periodontitis group, the initial mean PPD of 4.1 mm significantly decreased to 3.37 mm following NSPT (p < 0.0001; Figure 1B). This clinical recovery was further supported by significant reductions in BOP and the prevalence of plaque and calculus (p < 0.0001 for all; Figure S1 C,D).

**FIGURE 1.**
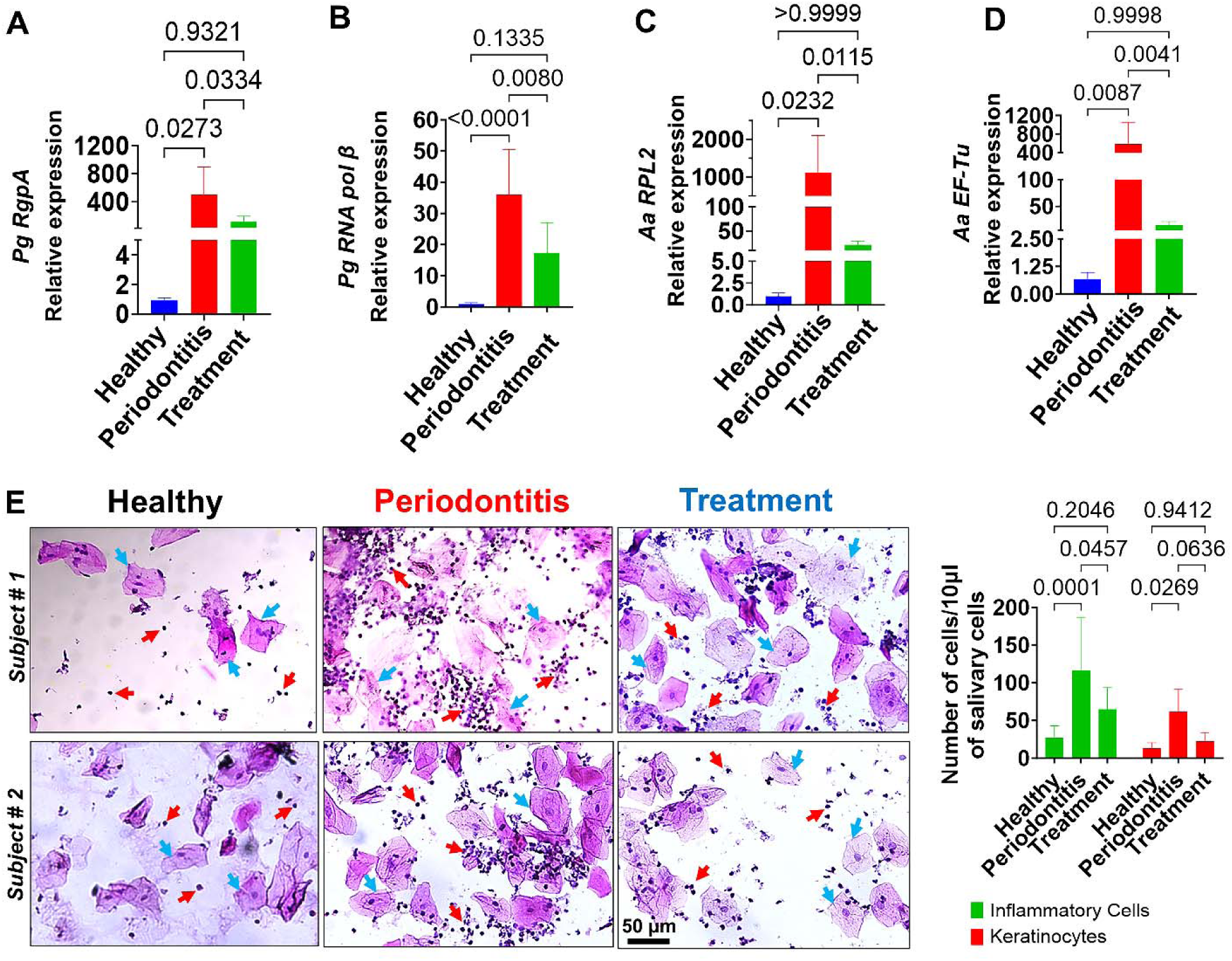
Salivary microbial and immune cell burden are elevated in periodontitis and respond to therapy. Salivary expression of (A) *P. gingivalis* RgpA gingipain, (B) *P. gingivalis* RNA pol β, (C) *A. actinomycetemcomitans* ribosomal protein L2 and (D) *A. actinomycetemcomitans* EF-Tu transcripts were evaluated in periodontally healthy subjects, untreated periodontitis (PD), and post–non-surgical periodontal therapy (NSPT) groups. (E) Assessment of salivary epithelial and immune cells abundance using Giemsa staining in periodontitis and post-NSPT subjects (left panel). Quantification of keratinocytes and immune cells in different groups (right panel). All analyses were conducted on saliva samples from periodontitis subjects before therapy (n = 18) and after therapy (n = 18) and compared with healthy controls (n = 12). Statistical analysis was performed using one-way ANOVA. Data are presented as mean ± SD. p < 0.05, p < 0.01, *p* < 0.001.

To assess the impact of NSPT on the oral microbiome, we quantified salivary bacterial-derived gene transcripts in diseased subjects at baseline and post-treatment. Based on prior RNA-seq profiling of gingival tissues, Pg- and Aa-encoded virulence factors were identified as highly expressed candidate transcripts (Uttamani et al., 2024). Compared with healthy controls, subjects with periodontitis showed significantly higher expression of RgpA (∼500-fold), RNA polymerase β (∼36-fold), RPL2 (∼1,100-fold), and EF-Tu (∼425-fold) (p<0.05; Figure 1A–D). Following NSPT, transcript levels of Pg RgpA and RNA polymerase β decreased to 109- and 18-fold (p<0.05), respectively. Similarly, Aa RPL2 and EF-Tu expression levels were reduced to 21- and 15-fold (p<0.05), respectively, compared with baseline. Despite this reduction, transcript levels in post-therapy samples remained above those observed in healthy controls.

Chronic periodontitis induces significant changes in the cellular composition of saliva, largely due to periodontal tissue destruction (Matsuoka et al., 2025). To determine whether reducing the periodontal bacterial burden influences salivary immune cell abundance, we performed salivary cytology using Giemsa staining in healthy, periodontitis, and post-therapy subjects. Microscopic examination revealed epithelial keratinocytes and immune cells across all groups. In healthy subjects, keratinocytes (blue arrows) and leukocytes (red arrows) were present in relatively balanced proportions, consistent with oral immune homeostasis (Figure 1C). Saliva from periodontitis subjects showed a marked increase in both the abundance and diversity of cellular populations, characterized by elevated leukocyte infiltration (115.85 ± 70.98) and increased epithelial cell shedding (61.22 ± 30.16), indicative of active inflammation associated with periodontal dysbiosis. Post-therapy samples exhibited substantial reductions in leukocyte (65 ± 28.56) and epithelial cell (22.55 ± 11.31) counts compared with diseased subjects, approaching levels observed in healthy controls (26.83 ± 15.76 leukocytes and 13.71 ± 6.76 epithelial cells). These findings suggest that lowering bacterial burden attenuates salivary cellularity (Figure 1E).

### 3.2 Elevated Myeloid and Lymphoid Populations in Saliva of Periodontitis Subjects

We first assessed whether salivary immune cell populations could discriminate between periodontal health, disease, and response to NSPT. Specifically, we quantified myeloid cell subsets (CD11b⁺CD14⁺HLA-DR⁺ and CD11b⁺CD11c⁺HLA-DR⁺) and lymphoid populations (CD4⁺ T cells, CD8⁺ T cells, and B cells). The total number of salivary CD11b⁺CD14⁺HLA-DR⁺ and CD11b⁺CD11c⁺HLA-DR⁺ myeloid cell populations was significantly elevated in diseased samples compared with healthy controls (22.94 ± 7.51% vs 11.78 ± 4.33%, p=0.007; 20.42 ± 7.85% vs 5.03 ± 2.38%, p<0.0001), respectively, consistent with previous reports demonstrating enhanced myeloid cell activation in periodontal inflammation (Figure 2A-D, Table 2). Similarly, lymphoid cell populations, including CD4⁺ T cells (29% vs 14%, p=0.002) and CD8⁺ T cells (0.88% vs 0.21%, p<0.015) were markedly elevated in the periodontitis group relative to healthy individuals (Figure 2E-G, Table 2). Following NSPT, a significant reduction in both myeloid and lymphoid cell populations was observed. The frequency of both CD11b⁺CD14⁺HLA-DR⁺ and CD11b⁺CD11c⁺HLA-DR⁺ myeloid cell subsets declined post therapy (22% vs 14%, p<0.0001; 20% vs 9%, p<0.003, Figure 2A-D, Table 2). Similarly, CD4⁺ T cells and CD8⁺ T cells demonstrated a substantial reduction following therapy (29% vs 19%, p<0.0003; 0.88% vs 0.39%, p<0.047, respectively, Figure 2E-G, Table 2). We also evaluated the proportions of CD4+ T and CD8+ T cells in gingiva from healthy and periodontitis subjects. Similar to salivary analysis, we noted a significant increase in gingival levels of CD4+ T cells and CD8+ T cells in the periodontitis cohort (25%, p = 0.0075 and 0.93%, p = 0.0029) compared to the healthy cohort (12% and 0.24%) (Figure 2h-J, Table 2). Consistent with prior reports (Dutzan et al.,2016, Malmqvist et al., 2024) and our preceding analysis, we observed a progressive decline in CD4+ and CD8+ T cells abundance levels in post-NSPT subjects compared to PD (Figure 2H-J, Table 2).

**FIGURE 2.**
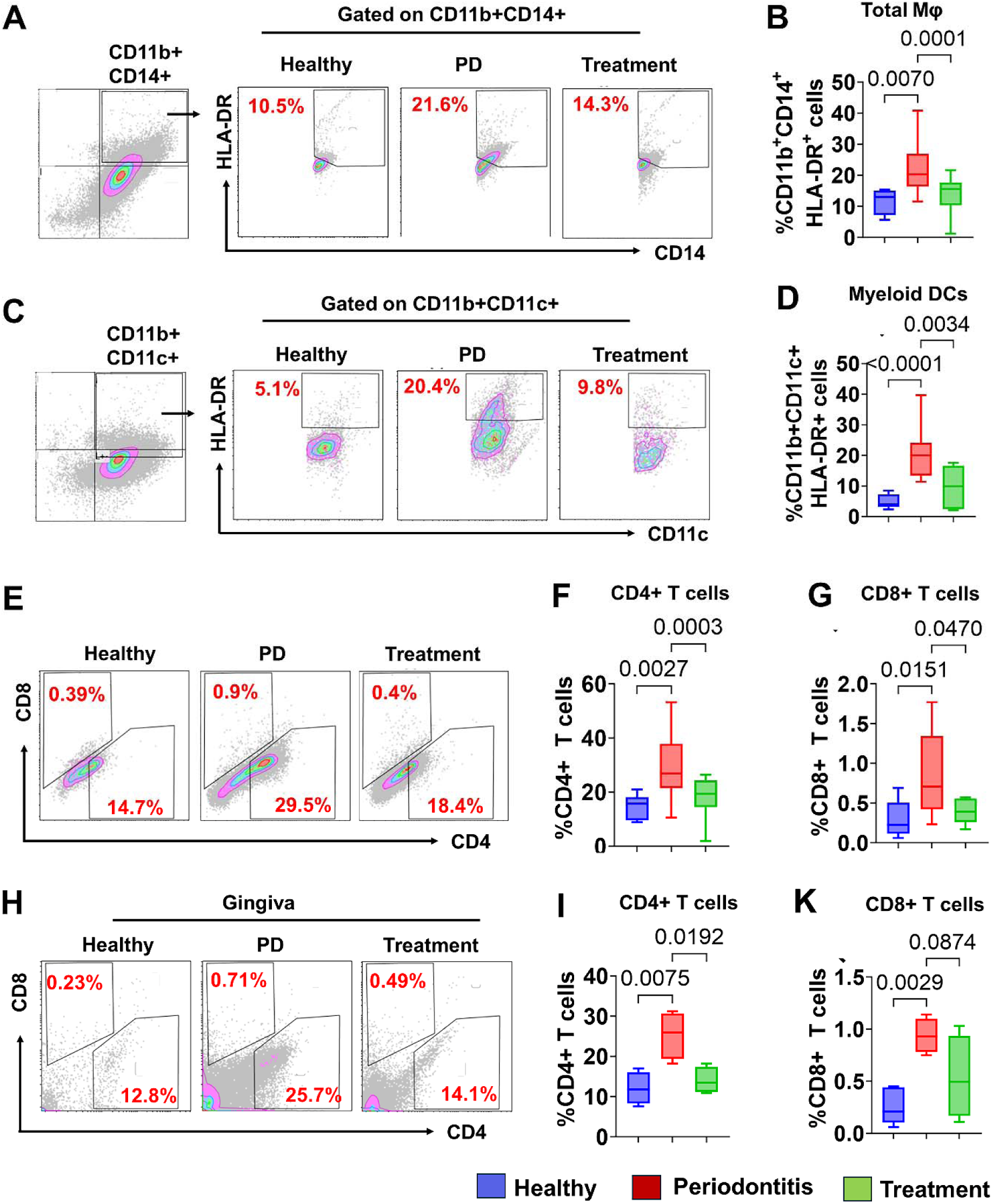
Periodontitis is associated with altered salivary myeloid and lymphoid immune cell frequencies that improve following therapy. Representative contour plots illustrate total macrophages (CD11b⁺CD14⁺HLA-DR⁺) (A), and dendritic cells (CD11b⁺CD11c⁺HLA-DR⁺) (C). CD4⁺ and CD8⁺ T-cell populations (E). Corresponding bar graphs depict total percentages of (B) macrophages, (D) dendritic cells, (F) CD4⁺ and (G) CD8⁺ T cells in all samples. All contour plots were generated using Kaluza software and bar graphs were produced using GraphPad. Statistical analysis was performed using one-way ANOVA. All analyses were conducted on saliva samples from periodontitis subjects before therapy (n = 18) and after therapy (n = 18), and compared with healthy controls (n = 12). Data are presented as mean ± standard deviation (SD), with statistical significance indicated as *p < 0.05 and **p < 0.01.

### 3.3 Salivary Pro-inflammatory Myeloid Cell Subsets are Increased in Periodontitis

To further characterize salivary myeloid cells, we assessed their polarization by measuring intracellular IFN-γ and IL-10 expressions. CD11b⁺CD14⁺HLA-DR⁺ cells expressing IFN-γ represent functional M1 Mφ, indicative of pro-inflammatory polarization capable of driving tissue destruction and activating adaptive immunity (Wang et al., 2023, Wu et al., 2025). Similarly, CD11b⁺CD11c⁺HLA-DR⁺IFN-γ⁺ reflect a second myeloid subset with strong pro-inflammatory potential (Collin et al., 2018, Li et al., 2025). Both myeloid cell subsets producing IFN-γ were significantly elevated in the saliva of periodontitis subjects compared with healthy individuals (CD11b⁺CD14⁺HLA-DR⁺IFN-γ⁺: 15% vs 7%; p=0.001; CD11b⁺CD11c⁺HLA-DR⁺IFN-γ⁺:17% vs 4%, p = 0.0009), (Figure 3A,B,E,F). Particularly, following NSPT, the frequency of these pro-inflammatory subsets decreased markedly (15% vs 10%, p = 0.026; 16% vs 11%, p < 0.024, respectively, Figure 3A,B,E,F; Table 2), mirroring reductions in adaptive lymphoid populations and reflecting improvements in clinical outcomes. These findings indicate that the abundance of IFN-γ⁺ myeloid cells in saliva not only correlates with disease severity but also effectively tracks therapeutic response, supporting their utility as a functional biomarker in periodontitis.

**FIGURE 3.**
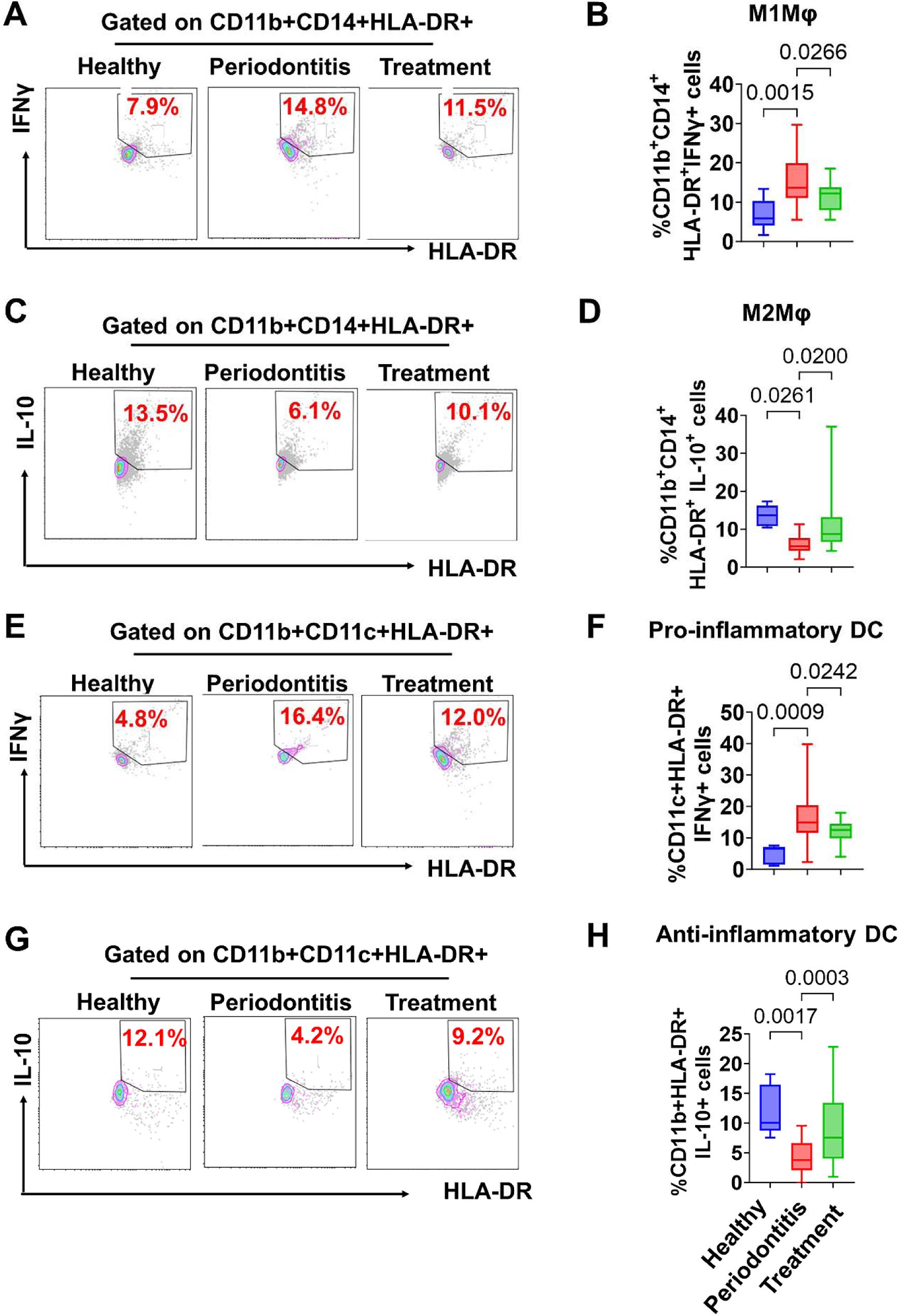
Periodontal therapy restores salivary macrophage and dendritic cell immune homeostasis in periodontitis. Representative contour plots show IFN-γ–producing pro-inflammatory macrophages (CD11b⁺CD14⁺HLA-DR⁺IFN-γ⁺; M1 macrophages) (A), IL-10–producing anti-inflammatory macrophages (CD11b⁺CD14⁺HLA-DR⁺IL-10⁺; M2 macrophages) (C), pro-inflammatory dendritic cells (CD11b⁺CD11c⁺HLA-DR⁺IFN-γ⁺) (E), and immunosuppressive dendritic cells (CD11b⁺CD11c⁺HLA-DR⁺IL-10⁺) (G). Corresponding bar graphs depict the frequencies of M1 macrophages (B), M2 macrophages (D), IFN-γ–producing dendritic cells (F), and IL-10–producing immunosuppressive dendritic cells (H). All contour plots were generated using Kaluza software, and bar graphs were produced using GraphPad. Statistical analysis was performed using one-way ANOVA. All analyses were conducted on saliva samples from periodontitis subjects before therapy (n = 18) and after therapy (n = 18), and compared with healthy controls (n = 12). Data are presented as mean ± standard deviation (SD), with statistical significance indicated as *p < 0.05 and **p < 0.01.

Next, we evaluated anti-inflammatory myeloid populations by measuring IL-10 expression. CD11b⁺CD14⁺HLA-DR⁺IL-10⁺ (Chen et al., 2023, Yun et al., 2025) and CD11b⁺CD11c⁺HLA-DR⁺IL-10⁺ cells (Boks et al., 2012, Comi et al., 2018) represent regulatory phenotypes that can counterbalance pro-inflammatory responses. The percentage of CD11b⁺CD14⁺HLA-DR⁺IL-10⁺ myeloid cells was significantly lower in periodontitis subjects compared with healthy controls (6% vs 13%, p = 0.026), levels of which increased post NSPT (6% vs 11%, p = 0.020). Furthermore, the levels of CD11b⁺CD11c⁺HLA-DR⁺IL-10⁺ cells were significantly lower in periodontitis subjects (∼4% vs 12%, p<0.001). Following NSPT, these populations showed a modest increase to 9%, suggesting partial restoration of anti-inflammatory signaling (Figure 3C,D,G,H; p<0.0005).

Overall, these findings indicate that the equilibrium between pro-inflammatory (IFN-γ⁺) and anti-inflammatory (IL-10⁺) myeloid subsets is dysregulated in periodontitis, favoring a sustained inflammatory environment, whereas NSPT partially restores this equilibrium.

### 3.4 Elevated Th1 and Th17 Cells in Saliva Corelate with Periodontal Disease Severity

T cell polarization is associated with PD pathogenesis and its resolution (Ebersole et al., 1994; Cardoso et al., 2009; Cavalla et al., 2022; Bi et al., 2019). To assess how T cell polarization is reflected in saliva of PD subjects, we characterized CD4⁺ T cell subsets, focusing on Th1, Th17, and Tr1 populations. We observed higher frequencies of Th1 and Th17 cells in periodontitis subjects compared with healthy controls. Following NSPT, both Th1 and Th17 populations were markedly reduced (Figure 4A-D), indicating that treatment dampens pathogenic T helper responses. To determine whether the systemic salivary profile accurately reflects localized tissue pathology, we validated the salivary immune phenotype within corresponding gingival biopsies. Flow cytometric analysis of gingival tissue revealed a CD4^+^IFNγ^+^ cell distribution pattern that closely paralleled our salivary observations. Compared to healthy gingiva (∼7%), significantly higher proportion of CD4^+^IFNγ^+^ T cells (∼20%, p = 0.03) was detected in the inflamed gingiva, followed by a reduction in the post-NSPT cohort (∼14%; p>0.05) (Figure 4G,H).

**FIGURE 4.**
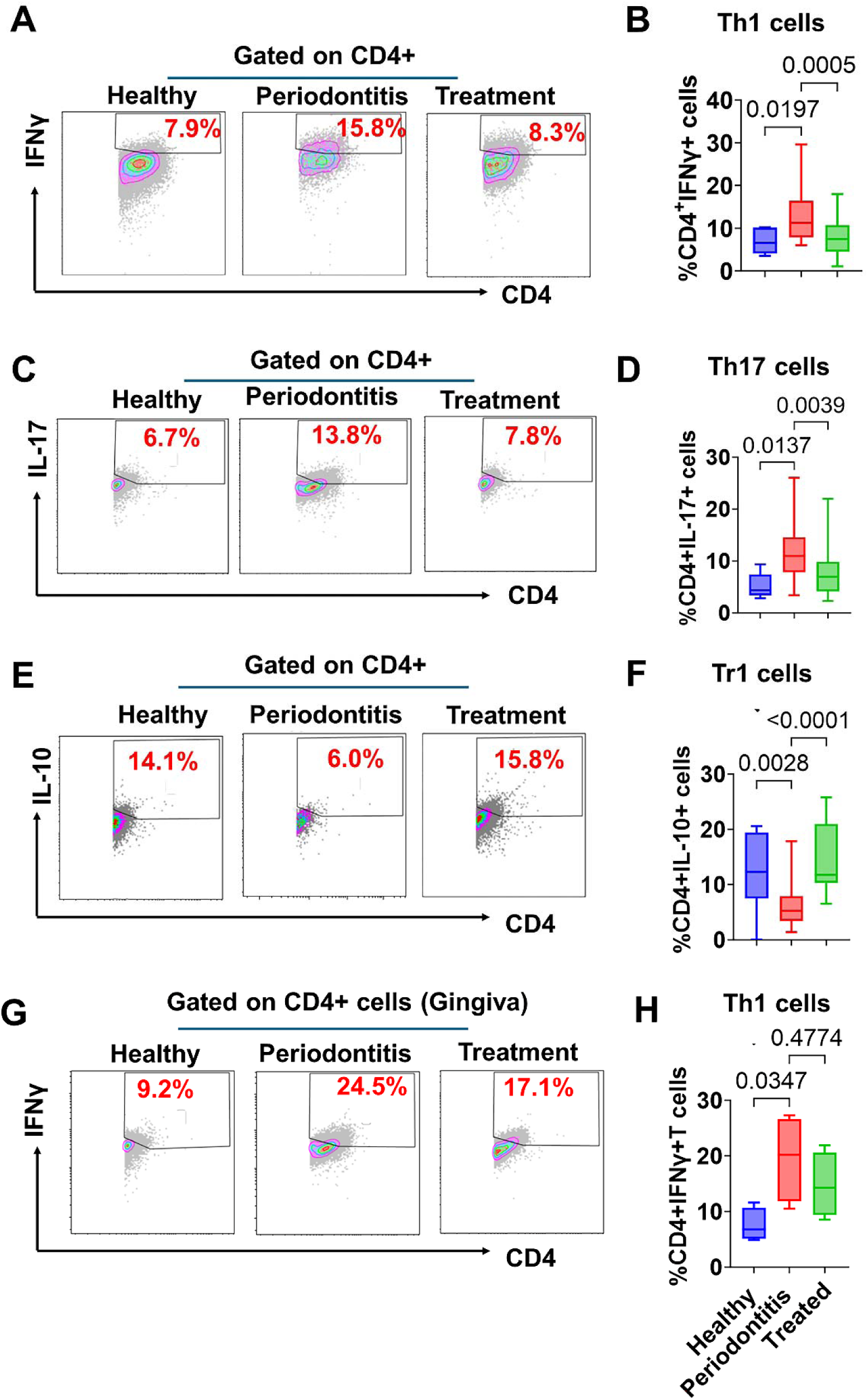
Periodontitis-associated salivary Th1 and Th17 responses improve after non-surgical periodontal therapy. Salivary profiles of Th1, Th17, and Tr1 immune responses in periodontitis subjects before and after therapy, compared with healthy controls. Representative contour plots illustrate Th1 (CD4+IFNγ+) (A), Th17(CD4+IL-17+) (C), and Tr1 (CD4+ IL-10+) (E) cell populations, with corresponding bar graphs showing Th1 (B), Th17 (D), and Tr1 (F) responses. Bar graphs represent the relative frequencies obtained by flow cytometric analysis of saliva samples from periodontitis subjects before therapy (n=18) and after therapy (n = 18), compared with healthy controls (n = 12). Statistical analysis was performed using Student’s t-test. Data are presented as mean ± standard error of the mean (SEM), with statistical significance indicated as *p < 0.05 and **p < 0.01

Regulatory Tr1 cells, which produce IL-10 and contribute to immune homeostasis, were evaluated to assess their potential role in restoring balance between pro-inflammatory T cell subsets. Tr1 cells were significantly lower in saliva from periodontitis subjects compared to healthy controls. Notably, numbers of Tr1 cells significantly increased after NSPT compared to periodontitis subjects suggesting partial re-establishment of immune regulation and equilibrium between pro-and anti-inflammatory T cell populations (Figure 4E,F, Table 2). Furthermore, evaluation of Tr1 response in the gingival tissue of pre-and post-NSPT subjects revealed a similar pattern as observed in the saliva. We observed a significant decline in the proportion of CD4^+^IL-10^+^ cells in the gingiva of periodontitis subjects compared to healthy gingival tissue, which was followed by a remarkable increase in post-NSPT subjects (Figure 4I,J, Table 2).

Together, these findings indicate that salivary Th1 and Th17 cells are co-existing contributors to periodontal inflammation and may serve as potential salivary biomarkers of disease activity and progression, while Tr1 cells may reflect therapy-mediated restoration of immunological balance.

### 3.5 Periodontitis Drives Salivary Accumulation of Pro-Inflammatory B Cells

B cells play a critical role in the pathogenesis of periodontitis by contributing to both host defense and tissue destruction through antibody production, cytokine secretion, and receptor activator of nuclear factor kappa-B ligand (RANKL)-mediated bone resorption (Figueredo et al., 2019; Han et al.; 2007, Taubman et al., 2007). In our periodontitis cohort, total CD19⁺ B cells were significantly increased compared with healthy controls (0.95% vs 0.39%; p = 0.042) and were markedly reduced following NSPT (0.24%; p = 0.0007) (Figure 5A,B), approaching levels observed in healthy individuals. To further delineate B cell function, we assessed pro-inflammatory B cell subsets defined as CD19⁺IFN-γ⁺ cells. Notably, this study demonstrates for the first time the presence of IFN-γ–producing B cells in saliva, which were significantly elevated in periodontitis subjects compared with healthy controls (21.3% vs 3.5%, p = 0.014), levels of which decreased following NSPT (8.9%, p = 0.141) (Figure 5C,D). These findings suggest that B cells may directly contribute to the pro-inflammatory milieu and amplification of adaptive immune responses in periodontitis.

**FIGURE 5.**
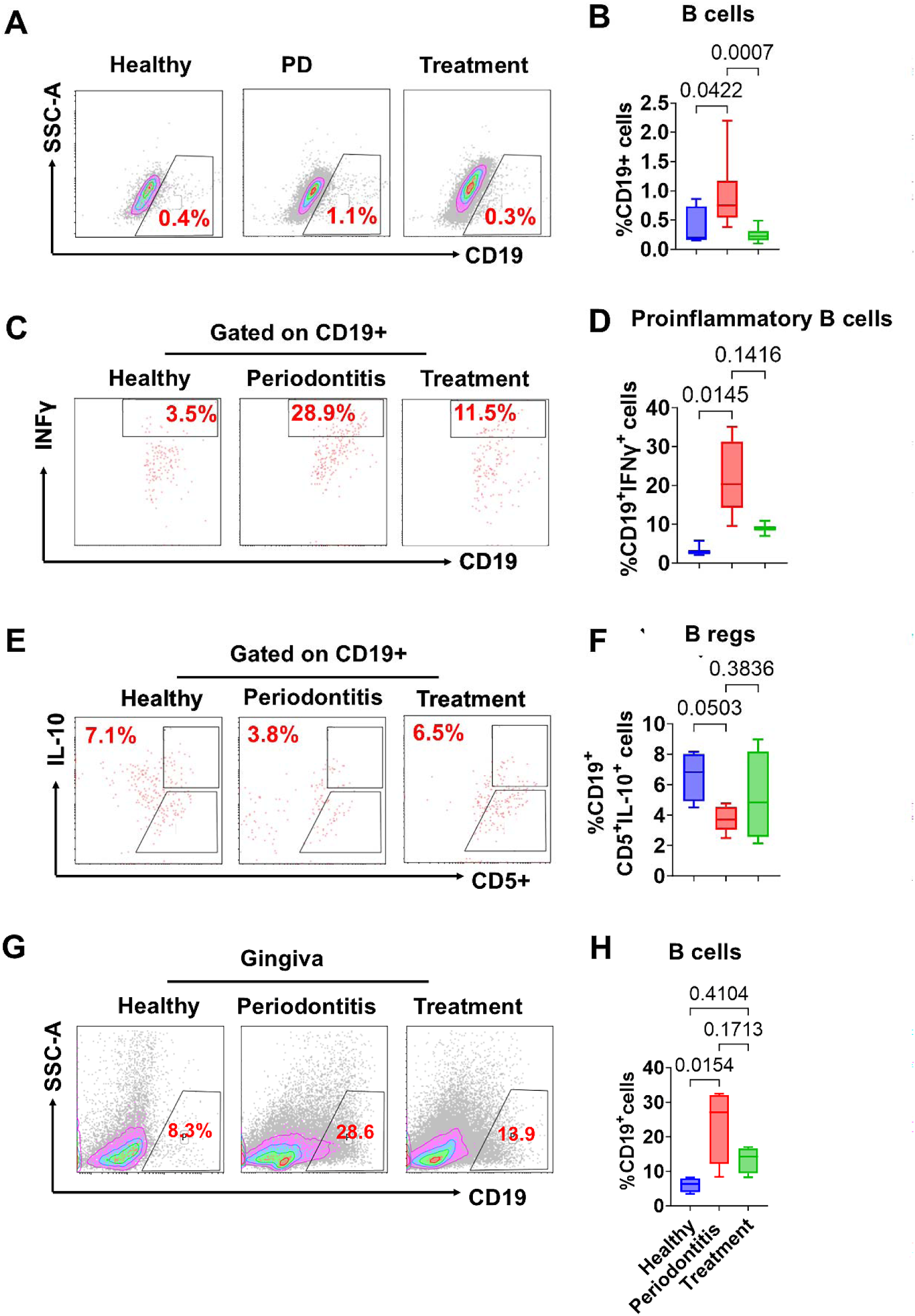
Periodontal therapy re-establishes salivary B-cell immune homeostasis by reducing IFN-γ–producing B cells and restoring regulatory B-cell frequencies. Representative contour plots show total B cells (CD19⁺) (A), IFN-γ–producing pro-inflammatory B cells (CD19⁺IFN-γ⁺) (C), and regulatory B cells (CD19⁺CD5⁺IL-10⁺) (E), with corresponding bar graphs presented in (B), (D), and (F), respectively. Bar graphs represent the relative frequencies of these populations as determined by flow cytometric analysis of saliva samples from periodontitis subjects before therapy (n = 18) and after therapy (n = 18), compared with healthy controls (n = 12). Statistical analysis was performed using Student’s t-test. Data are presented as mean ± standard error of the mean (SEM) from three independent experiments performed in triplicate, with statistical significance indicated as *p < 0.05 and **p < 0.01.

Conversely, regulatory B cells (Bregs), identified as CD19⁺CD5⁺IL-10⁺ cells, were significantly reduced in subjects with periodontitis compared with healthy controls (3.73 ± 0.85% vs. 7.13 ± 1.4%; p < 0.05). Following NSPT, Breg levels increased to 5.19 ± 2.93%, although this change did not reach statistical significance (p = 0.38) (Figure 5E,F). This trend suggests a partial restoration of anti-inflammatory immune function after therapy. A similar pattern was observed in gingival tissues. Periodontitis patients showed a significant increase in CD19⁺ B-cell infiltration in gingival tissues (3.1%) compared with healthy gingiva (0.63%; p = 0.009). Following NSPT, the B-cell population was markedly reduced to 1.21% (p = 0.04) (Figure 5G,H), consistent with the changes observed in saliva samples.

These results demonstrate that B cell populations in periodontitis are skewed toward a pro-inflammatory phenotype, with increased IFN-γ–producing B cells and reduced regulatory B cells. NSPT partially restores this imbalance, suggesting that B cell functional equilibrium is disrupted in the pro-inflammatory microenvironment of periodontitis and rebalanced following therapy.

## 4 Discussion

The present study identifies saliva as a clinically informative and mechanistically relevant biospecimen for interrogating immune cell populations underlying periodontitis and its response to therapy. Using multidimensional immune phenotyping, we demonstrate that saliva captures alterations in immune-cell composition associated with active disease and therapeutic resolution. To our knowledge, this is the first study to provide comprehensive salivary immunophenotyping across periodontal health, active disease, and post-therapeutic resolution. Importantly, comparative immune profiling of gingival biopsy specimens demonstrated substantial concordance between salivary and tissue-level immune signatures, indicating that salivary immune profiles reflect the gingival immune microenvironment. These findings suggest that saliva harbors a dynamic immunological compartment capable of capturing ongoing microbial and periodontal immune activity.

A major strength of this study lies in the depth of salivary immune profiling. Although prior investigations have focused largely on soluble inflammatory mediators, comprehensive cellular immunophenotyping in saliva remains limited. Here, we demonstrate broad expansion of both innate and adaptive immune populations in periodontitis, including myeloid cells, T-cell subsets, and B-cell populations, all of which were partially normalized following NSPT in parallel with reduced microbial burden and clinical improvement.

At the level of innate immunity, periodontitis was characterized by a pronounced shift toward a pro-inflammatory myeloid phenotype, with increased IFN-γ–producing (M1-like) cells and reduced IL-10–producing (M2-like) regulatory populations. NSPT partially restored this imbalance, suggesting that periodontal therapy modulates not only microbial burden but also myeloid immune programming. These findings are consistent with current understanding that unresolved inflammatory myeloid responses contribute directly to periodontal tissue destruction and impaired inflammatory resolution (Hajishengallis et al., 2020; Uttamani et al., 2024; Baima et al., 2025). Restoration of IL-10–associated regulatory pathways after therapy may represent an essential component of tissue stabilization and immune homeostasis.

Within the adaptive immune compartment, our findings extend beyond the traditional Th17-centric framework of periodontitis. Although Th17 cells were significantly elevated, we also identified concurrent expansion of Th1 cells, indicating coordinated activation of multiple effector T-cell pathways. The coexistence of IFN-γ– and IL-17–producing T cells likely reflects chronic stimulation by a dysbiotic microbial ecosystem capable of engaging diverse inflammatory pathways simultaneously (Gaffen et al., 2008; Dutzan et al., 2018). Importantly, both Th1 and Th17 populations declined following NSPT, whereas regulatory Tr1 cells showed modest recovery, consistent with partial restoration of immune homeostasis. These observations suggest that disease progression is driven not by a single dominant T-cell lineage, but by broader dysregulation of effector and regulatory immune networks.

This study provides novel insights into B-cell biology in periodontitis, particularly within saliva. Although B cells are established contributors to periodontal pathology through antibody production and RANKL-mediated osteoclastogenesis (Settem et al., 2021), their functional polarization in saliva has not been characterized. We demonstrate, for the first time, the presence of IFN-γ–producing B cells in saliva and show that these pro-inflammatory B-cell populations are significantly expanded in periodontitis. Conversely, regulatory B cells (CD19⁺CD5⁺IL-10⁺) were diminished during disease. Both cell types were partially restored after therapy. These findings indicate that B-cell responses in periodontitis extend beyond humoral immunity and involve active participation in inflammatory immune orchestration. The identification of distinct salivary B-cell subsets adds a previously unrecognized dimension to periodontal immunopathology and highlights their potential as biomarkers and therapeutic targets.

These findings have important translational implications. Unlike conventional periodontal diagnostics, which primarily reflect past tissue destruction, salivary profiling provides a minimally invasive, real-time assessment of microbial and immune activity. Compared with gingival biopsies, saliva offers rapid, reproducible, and scalable access to immune cells for longitudinal monitoring. Persistence of immune and microbial signatures despite clinical improvement suggests that residual inflammation and dysbiosis may remain undetected by conventional clinical measures. The ability to simultaneously quantify bacterial transcripts and immune-cell phenotypes from a single biospecimen creates opportunities for integrated salivary biomarkers to identify active diseases, predict treatment response, and monitor recurrence. Furthermore, the strong concordance between gingival and salivary immune signatures supports saliva as an accessible surrogate of the periodontal immune environment.

This study has limitations. The sample size was modest, potentially limiting statistical power for less abundant immune subsets. The observational design precludes definitive conclusions regarding causal relationships between microbial dysbiosis and immune-cell polarization. In addition, bacterial analyses focused on selected periodontal pathogens and may not fully capture the oral microbiome’s complexity. Future studies integrating larger longitudinal cohorts, microbiome-wide analyses, and single-cell immune profiling will be important to define mechanistic relationships between microbial persistence, immune remodeling, and clinical outcomes. Nevertheless, this study establishes saliva as a clinically tractable platform for studying periodontal immunopathology and provides a foundation for precision immunodiagnostic approaches in periodontitis.

## Conclusion

This study provides one of the most comprehensive characterizations of salivary immune subsets to date, including previously underappreciated B-cell functional phenotypes across health, disease, and post-treatment states. Collectively, these data support saliva as a scalable, non-invasive platform for precision diagnostics and longitudinal monitoring in periodontitis.

## Author Contributions

All authors have made substantial contributions to conception, design of the study and given final approval of the version to be published. MT, KC and JG were calibrated and performed clinical data measurements, as well as obtained biomaterials from subjects. RAN, MT, KC, JG, SE, LD, MLS, JLS, SN, and ARN were involved in data collection and data analysis. RAN, JLS, SN, and ARN were involved in data interpretation, and drafting the manuscript.

## Acknowledgements

The authors gratefully acknowledge the staff of the University of Illinois Chicago College of Dentistry and the study participants for their valuable contributions to this research.

## Data Availability Statement

The data that support the findings of this study are available from the corresponding author upon reasonable request.

## Ethics Statement

The present study was evaluated by the ethics committee of the University of Illinois Chicago. Written informed consent was obtained from all participants prior to their enrolment in the study.

## References

1. Könönen E, Gursoy M, Gursoy UK. Periodontitis: A multifaceted disease of tooth-supporting tissues. J Clin Med. 2019;8(8):1135.

2. Gasner NS, Schure RS. Periodontal disease. In: StatPearls. Treasure Island, FL: StatPearls Publishing; 2024. Available from: http://www.ncbi.nlm.nih.gov/books/NBK554590/ (accessed 19 April 2024).

3. Kinane DF, Stathopoulou PG, Papapanou PN. Periodontal diseases. Nat Rev Dis Primers. 2017;3:17038.

4. Wang Y, Zhao X, Yu Y. The burden of severe periodontitis in the United States. J Am Dent Assoc. 2025;157:123–132.e16.

5. O’Dwyer MC, Furgal A, Furst W, Ramakrishnan M, Capizzano N, Sen A, et al. The prevalence of periodontitis among US adults with multimorbidity using NHANES data 2011–2014. J Am Board Fam Med. 2023;36(2):313–324.

6. Sanz-Martín I, Cha JK, Yoon SW, Sanz-Sánchez I, Jung UW. Long-term assessment of periodontal disease progression after surgical or non-surgical treatment: A systematic review. J Periodontal Implant Sci. 2019;49(2):60–75.

7. Loos BG, Needleman I. Endpoints of active periodontal therapy. J Clin Periodontol. 2020;47(Suppl 22):61–71.

8. Tonetti MS, Greenwell H, Kornman KS. Staging and grading of periodontitis: Framework and proposal of a new classification and case definition. J Periodontol. 2018;89(Suppl 1):S159–S172.

9. Liukkonen J, Gürsoy UK, Könönen E, et al. Salivary biomarkers in association with periodontal parameters and the periodontitis risk haplotype. Innate Immun. 2018;24(7):439–447.

10. Surdu A, Foia LG, Luchian I, Trifan D, Tatarciuc MS, Scutariu MM, et al. Saliva as a diagnostic tool for systemic diseases: A narrative review. Medicina (Kaunas). 2025;61(2):243.

11. Ji S, Choi Y. Point-of-care diagnosis of periodontitis using saliva: Technically feasible but still a challenge. Front Cell Infect Microbiol. 2015;5:65.

12. Isho B, Abe KT, Zuo M, Jamal AJ, Rathod B, Wang JH, et al. Persistence of serum and saliva antibody responses to SARS-CoV-2 spike antigens in COVID-19 patients. Sci Immunol. 2020;5(52):eabe5511.

13. Miller CS, Foley JD, Bailey AL, Campell CL, Humphries RL, Christodoulides N, et al. Current developments in salivary diagnostics. Biomark Med. 2010;4(1):171–189.

14. Mehrnia N, Van Dyke TE. Microbial dysbiosis and immune dysregulation in periodontitis and peri-implantitis. Front Cell Infect Microbiol. 2026;15:1678163.

15. Baima G, Arce M, Romandini M, Van Dyke T. Inflammatory and immunological basis of periodontal diseases. J Periodontal Res. 2025. doi:10.xxxx/jre.xxxxx.

16. Hajishengallis G. Immunomicrobial pathogenesis of periodontitis: Keystones, pathobionts, and host response. Trends Immunol. 2014;35(1):3–11.

17. Hajishengallis G, Lamont RJ. Polymicrobial communities in periodontal disease: Their quasi-organismal nature and dialogue with the host. Periodontol 2000. 2021;86:210–230.

18. Matsuoka M, Soria SA, Pires JR, Sant’Ana ACP, Freire M. Natural and induced immune responses in oral cavity and saliva. BMC Immunol. 2025;26(1):34.

19. Vidović A, Vidović Juras D, Vučićević Boras V, Lukač J, Grubišić-Ilić M, Rak D, et al. Determination of leucocyte subsets in human saliva by flow cytometry. Arch Oral Biol. 2012;57:577–583.

20. Choudhury SN, Novotny M, Aevermann BD, et al. A protocol for revealing oral neutrophil heterogeneity by single-cell immune profiling in human saliva. Protoc Exch. 2020. doi:10.21203/rs.3.pex-953/v2.

21. Theda C, Hwang SH, Czajko A, et al. Quantitation of the cellular content of saliva and buccal swab samples. Sci Rep. 2018;8:6944.

22. Hajishengallis G, Chavakis T. Local and systemic mechanisms linking periodontal disease and inflammatory comorbidities. Nat Rev Immunol. 2021;21:426–440.

23. Uttamani JR, Naqvi AR, Estepa AMV, Kulkarni V, Brambila MF, Martínez G, et al. Downregulation of miRNA-26 in chronic periodontitis interferes with innate immune responses and cell migration by targeting phospholipase C beta 1. J Clin Periodontol. 2023;50(1):102–113.

24. Valverde A, Naqvi RA, Chen Y, Moshaverinia A, George A, Shukla D, et al. Herpes simplex virus-1 exploits inflammation to infect periodontal stem cells and disrupt lineage commitment. J Periodontal Res. 2025;60(12):1265–1279.

25. Naqvi RA, Valverde A, Nares S, Van Dyke TE, Naqvi AR. Myeloid-derived suppressor cells mitigate inflammation in periodontal disease. bioRxiv. 2024. doi:10.1101/2024.12.21.629927.

26. Teles FRF, Chandrasekaran G, Martin L, Patel M, Kallan MJ, Furquim C, et al. Salivary and serum inflammatory biomarkers during periodontitis progression and after treatment. J Clin Periodontol. 2024;51(12):1619–1631.

27. Uttamani JR, Kulkarni V, Valverde A, Naqvi RA, Van Dyke T, Nares S, et al. Dynamic changes in macrophage polarization during the resolution phase of periodontal disease. Immun Inflamm Dis. 2024;12(10):e70044.

28. Valverde A, Capistrano K, Naqvi RA, Elshourbagy S, Etminan S, Sandoval G, et al. Periodontitis primes the oral microenvironment for PS-dependent non-canonical entry pathways linked to SARS-CoV-2 susceptibility. Res Sq [Preprint]. 2026.

29. Parisi L, Gini E, Baci D, Tremolati M, Fanuli M, Bassani B, et al. Macrophage polarization in chronic inflammatory diseases: Killers or builders? J Immunol Res. 2018;2018:8917804.

30. Malmqvist S, Clark R, Johannsen G, Johannsen A, Boström EA, Lira-Junior R. Immune cell composition and inflammatory profile of human peri-implantitis and periodontitis lesions. Clin Exp Immunol. 2024;217(2):173–182.

31. Dutzan N, Konkel JE, Greenwell-Wild T, Moutsopoulos NM. Characterization of the human immune cell network at the gingival barrier. Mucosal Immunol. 2016;9(5):1163–1172.

32. Wu S, Mao Y, Hu L, Ji H, Liu X, Ma G, et al. Six interferon-stimulated genes as biomarkers of M1 macrophage polarization in psoriasis. J Inflamm Res. 2025;18:13835–13853.

33. Wang K, Mao T, Lu X, et al. A potential therapeutic approach for ulcerative colitis: Targeted regulation of macrophage polarization through phytochemicals. Front Immunol. 2023;14:1155077.

34. Collin M, Bigley V. Human dendritic cell subsets: An update. Immunology. 2018;154(1):3–20.

35. Li W, Yu C, Zhang X, Gu Y, He X, Xu R, et al. Dendritic cells: Understanding ontogeny, subsets, functions, and their clinical applications. Mol Biomed. 2025;6(1):62.

36. Chen S, Saeed AF, Liu Q, et al. Macrophages in immunoregulation and therapeutics. Signal Transduct Target Ther. 2023;8:207.

37. Yu B, Wang X, Zheng Y, Wang W, Cheng X, Cao Y, et al. M2 macrophages promote IL-10+ B-cell production and alleviate asthma in mice. Immunother Adv. 2025;5(1):ltaf007.

38. Boks MA, Kager-Groenland JR, Haasjes MS, Zwaginga JJ, van Ham SM, ten Brinke A. IL-10-generated tolerogenic dendritic cells are optimal for functional regulatory T-cell induction: A comparative study of human clinical-applicable DC. Clin Immunol. 2012;142(3):332–342.

39. Comi M, Amodio G, Gregori S. Interleukin-10-producing DC-10 is a unique tool to promote tolerance via antigen-specific T regulatory type 1 cells. Front Immunol. 2018;9:682.

40. Sun X, Gao J, Meng X, Lu X, Zhang L, Chen R. Polarized macrophages in periodontitis: Characteristics, function, and molecular signaling. Front Immunol. 2021;12:763334.

41. Cavalla F, Hernández M. Polarization profiles of T lymphocytes and macrophage responses in periodontitis. Adv Exp Med Biol. 2022;1373:195–208.

42. Bi CS, Sun LJ, Qu HL, Chen F, Tian BM, Chen FM. The relationship between T-helper cell polarization and the RANKL/OPG ratio in gingival tissues from chronic periodontitis patients. Clin Exp Dent Res. 2019;5(4):377–388.

43. Ebersole JL, Taubman MA. The protective nature of host responses in periodontal diseases. Periodontol 2000. 1994;5:112–141.

44. Cardoso CR, Garlet GP, Crippa GE, Rosa AL, Júnior WM, Rossi MA, et al. Evidence of the presence of T helper type 17 cells in chronic lesions of human periodontal disease. Oral Microbiol Immunol. 2009;24(1):1–6.

45. Figueredo CM, Lira-Junior R, Love RM. T and B cells in periodontal disease: New functions in a complex scenario. Int J Mol Sci. 2019;20(16):3949.

46. Han X, Kawai T, Taubman MA. Interference with immune-cell-mediated bone resorption in periodontal disease. Periodontol 2000. 2007;45:76–94.

47. Taubman MA, Kawai T, Han X. The new concept of periodontal disease pathogenesis requires new and novel therapeutic strategies. J Clin Periodontol. 2007;34:367–369.

48. Johnston W, Rosier BT, Artacho A, Paterson M, Piela K, Delaney C, et al. Mechanical biofilm disruption causes microbial and immunological shifts in periodontitis patients. Sci Rep. 2021;11(1):9796.

49. Giannobile WV, Beikler T, Kinney JS, Ramseier CA, Morelli T, Wong DT. Saliva as a diagnostic tool for periodontal disease: Current state and future directions. Periodontol 2000. 2009;50:52–64.

50. Hajishengallis G, Chavakis T, Lambris JD. Current understanding of periodontal disease pathogenesis and targets for host-modulation therapy. Periodontol 2000. 2020;84(1):14–34.

51. Dutzan N, Kajikawa T, Abusleme L, Greenwell-Wild T, Zuazo CE, Ikeuchi T, et al. A dysbiotic microbiome triggers TH17 cells to mediate oral mucosal immunopathology in mice and humans. Sci Transl Med. 2018;10(463):eaat0797.

52. Gaffen SL, Hajishengallis G. A new inflammatory cytokine on the block: Re-thinking periodontal disease and the Th1/Th2 paradigm in the context of Th17 cells and IL-17. J Dent Res. 2008;87(9):817–828.

53. Settem RP, Honma K, Chinthamani S, Kawai T, Sharma A. B-cell RANKL contributes to pathogen-induced alveolar bone loss in an experimental periodontitis mouse model. Front Physiol. 2021;12:722859.

