## Supplemental Figures for "Longitudinal Salivary Immunophenotyping Reveals Distinct Cellular Signatures of Periodontal Disease Activity and Resolution"

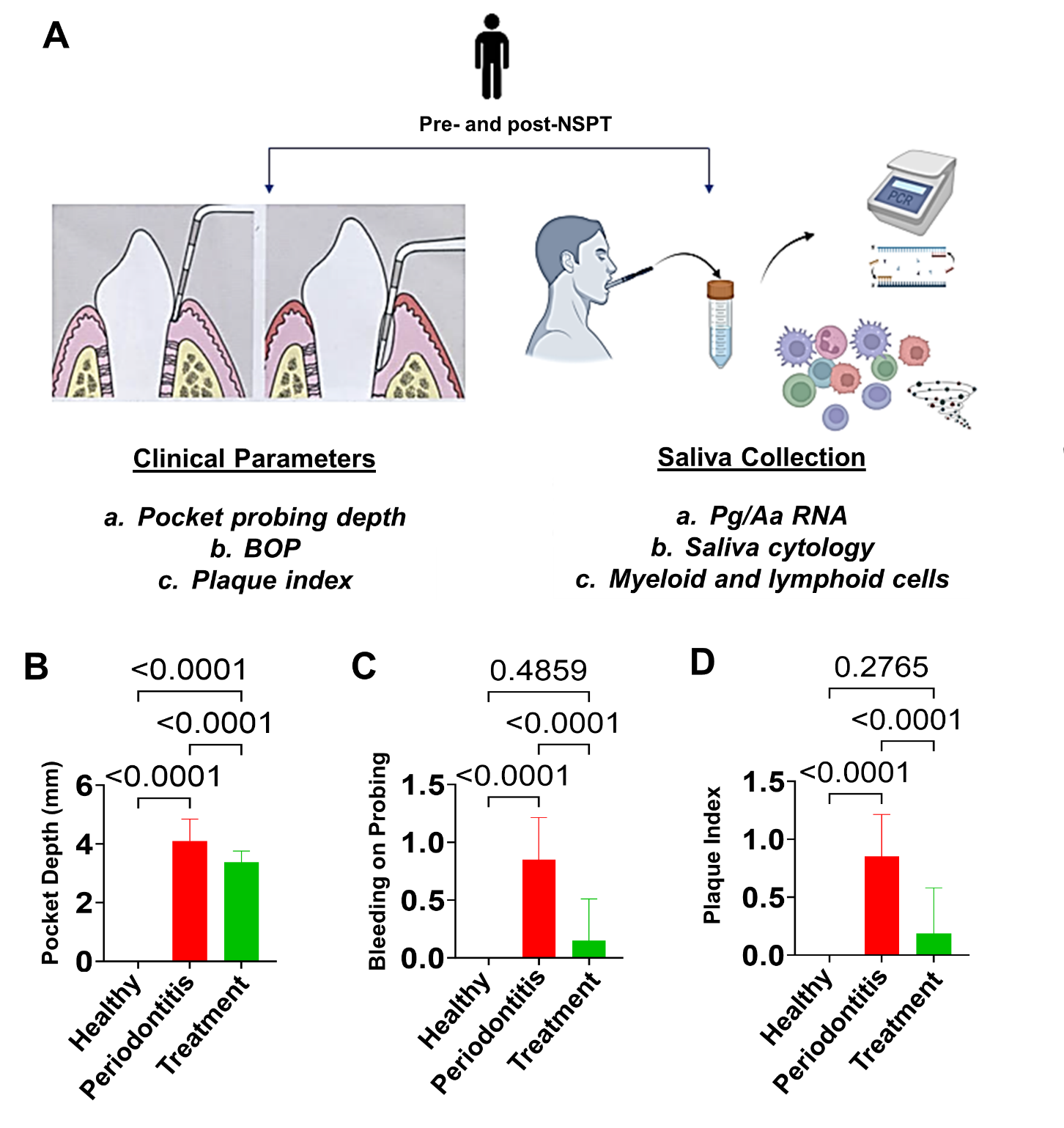
**FIGURE S1.** Study workflow and clinical periodontal parameters. (A) Schematic overview of the clinical examination and saliva collection protocol. Periodontal parameters, including probing depth (PPD), bleeding on probing (BOP), and plaque/calculus accumulation, were recorded, followed by collection of unstimulated whole saliva samples. Saliva samples were subsequently processed for downstream analyses, including immune cell immunophenotyping and molecular assessment of bacterial gene expression. Clinical periodontal parameters included probing depth (B), bleeding on probing (C), and plaque index (D).
